# Multiscale analysis of structurally conserved motifs

**DOI:** 10.1101/379768

**Authors:** F. Cazals, R. Tetley

## Abstract

This work develops a generic framework to perform a multiscale structural analysis of two structures (homologous proteins, conformations) undergoing conformational changes. Practically, given a seed structural alignment, we identify structural motifs with a hierarchical structure, characterized by three unique properties. First, the hierarchical structure sheds light on the trade-off between size and flexibility. Second, motifs can be combined to perform an overall comparison of the input structures in terms of combined RMSD - an improvement over the classical least RMSD. Third, motifs can be used to seed iterative aligners, and to design hybrid sequence-structure profile HMM characterizing protein families.

From the methods standpoint, our framework is reminiscent from the bootstrap and combines concepts from rigidity analysis (distance difference matrices), graph theory, computational geometry (space filling diagrams), and topology (topological persistence).

On challenging cases (class II fusion proteins, flexible molecules) we illustrate the ability of our tools to localize conformational changes, shedding light of commonalities of structures which would otherwise appear as radically different.

Our tools are available within the Structural Bioinformatics Library (http://sbl.inria.fr). We anticipate that they will be of interest to perform structural comparisons at large, and for remote homology detection.

## 1 Introduction

### 1.1 Structural alignment methods

The structure - function paradigm stipulates that the structure and the dynamics of proteins account for their function. Providing a structure based understanding of thermodynamic and kinetic molecular properties is however especially challenging, due to the vast array of spatial and temporal time-scales involved [1]. Depending on their focus, structure based studies are typically used to investigate two broad classes of problems. The first class is the analysis of (homologous) protein structures, a key endeavor for protein function annotation [2]. In this realm, finding structural alignments actually requires solving two problems at once: first, identifying the sub-sequences matching one-another; second, optimizing some criterion qualifying the geometry of the alignment. We note that these two problems are easy when solved independently: finding the alignment for fixed positions is amenable to dynamic programming; finding the optimal superimposition given the alignment is the classical rigid superimposition problem. The second class is the analysis of thermodynamics and dynamics from simulations (molecular dynamics, Monte Carlo sampling and their generalizations). In this setting, it is indeed well known that reliable thermodynamic and kinetic estimates requires accurate clustering algorithms to group conformations easily interconvertible into one another [3].

For these two classes of problems, a variety of structural alignment methods have been developed, which may be classified using their three main ingredients [4], namely the molecular representation used, the associated scoring function, and the optimization algorithm run.

- Classical representations favor geometric or topological features. The former are based on cartesian coordinates and/or internal distances. The latter are based on graphs coding geometric and/or topological properties. A popular class of graphs are contact maps, namely graphs whose edges code the spatial proximity between two amino-acids (a.a). Contact maps may be defined from pairs of *C_α_* which are within a distance threshold [5], or using physical contacts between a.a., such as those obtained from Voronoi models [6, 7]. We note that graph based representations tend to produce flexible alignments [5], as the network of contacts may be locally conserved yet globally geometrically distorted.
- Representations naturally call for specific scores. Geometric representations are typically assessed using the root mean square deviation of internal distances (dRMSD) [8, 9], the coordinate (least) root mean square deviations (lRMSD), or variants [10, 11]. More recently, geometric scores assessing the *compatibility* of backbone fragments using local frames have also been introduced [12]. Topological representations typically call for scores based on conserved contacts.
- Finally, optimization algorithms used are especially important when it comes to accommodate time execution constraints, and aligners may also be classified by the hardness of the problem solved. On the one hand, selected aligners solve easy i.e. polynomial time solvable problems. Approximate aligners, be they iterative [13, 14, 12] or not [10] fall in this category. On the other hand, search for optimal alignments in terms of contacts generally tackle NP-hard (and hard to approximate) optimization problems [15, 16], which in turns calls for approximate solutions when it comes to handling large scale comparisons [17].

As seen from this mini-review, the problem of performing structural alignments triggered developments in complementary directions, and pairwise comparisons have evidenced that alignments yielded by individual methods may indeed differ considerably [18]. This state of affairs prompted the development of tools easing the comparison of pairwise aligners [19].

### 1.2 Contributions

Consider two structures, typically two polypeptide chains, for which we wish to identify structurally conserved motifs. The structures may represent the same conformation of a given molecule, or the structures of two (homologous) polypeptide chains.

Our work is motivated by a refined analysis of flexible alignments, a delicate problem since flexibility is inherently related to scale-smaller regions tend to be more rigid. Consider a seed structural alignment. In order to perform a multiscale analysis of structural motifs, we are guided by the following principles:

- A motif is defined by an alignment between two sets of a.a. of the same size.
- Motifs must inherently exhibit a multiscale structure, trading size for structural conservation, and capturing the various structural conservation scales that may be present in the seed alignment.
- At any scale, motifs must be amenable to independent structural comparisons (using say lRMSD), as these lRMSD may be combined to assess more global deformations.

To meet these requirements, we make several contributions:

1. **A generic framework for multiscale structural alignments.** First, we introduce a generic bootstrap framework to report structurally conserved motifs, based on (i) a seed alignment, and (ii) a topological analysis of so-called *filtrations* coding conserved distances in the structures. We provide four instantiations obtained by combining two seed alignment methods (`Kpax` [12], `Apurva` [17]) and two *filtrations*. The rationale for using two seed alignment is as follows: `Apurva` is a structural aligner based on contact maps, favoring long and *flexible* alignments [17]; `Kpax` is a structural aligner based on a geometric representation of the backbone, favoring a geometric measure known as the G-score [12]. In a nutshell, a filtration is a sequence of nested topological spaces [20], which allows investigating a property, structural conservation in our case, at multiple scales. Practically, we use two filtrations. The first one, defined from distance difference matrices [8], favors distance conservation regardless of connectivity; the second one, defined from space filling diagrams [21], favors spatial connectivity.
2. **Dissecting a seed alignment: motifs and multiscale analysis.** We introduce an assessment of motifs based on Hasse diagrams to handle the multiscale nature of motifs, and Pareto fronts defined in the space (lRMSD, alignment size) of motifs. Using the latter, we investigate in particular the merits of the two filtrations used.
3. **Motifs to qualify flexible deformations via combined lRMSD.** We show that the optimal rigid motions associated to our motifs leverage the combined lRMSD, a notion of *flexible* lRMSD mixing local (motif based) lRMSD into a global score assessing a global deformation [22].
4. **Motifs to seed iterative alignments.** Iterative aligners consist in iteratively finding the alignment for fixed positions of the conformations and finding optimal superimposition given the alignment [14, 12]. We show that our motifs can be used to seed iterative structural alignments, yielding alignments outperforming those of the seed aligners (`Apurva`, `Kpax`).
5. **Software.** All our methods are made available from the SBL (http://sbl.inria.fr).

#### Validation: test-set

We use two datasets. The first one is a dataset of remote homologous class II fusion proteins. The second one is a dataset of ten challenging cases for flexible aligners.

Two important comments are in order. First, our generic method is not *stricto sensu* a direct competitor of flexible aligners. Indeed, a key goal is to dissect a seed alignment into motifs, so as to capture the various structural conservation scales it may contain. Second, our approach bears two major differences with the flexible alignment methods FATCAT [23] and the phenotypic plasticity measure (PPM) [24]. Both methods require the prior identification of rigid blocks. In [23], a flexible alignment is obtained using dynamic programming, by interleaving the blocks (called aligned fragment pairs) with connexions defined by extensions, gaps or twists. In [24], a flexible alignment is then defined by a spanning tree connecting blocks, and the associated PPM score measures the similarity between nodes (blocks) and edges (connexions between blocks). Once all possible blocks and topological edges connecting them have been computed, the optimal alignment is obtained from the A* algorithm. As opposed to these two methods, we do not rely on precomputed blocks, but identify them in a multiscale fashion; also, we perform a multiscale analysis of motifs, which are qualified in terms of Hasse diagrams and Pareto envelopes.

## 2 Method: extracting structural motifs

Our method to detect structural motifs mixes ingredients from structural alignments, graph theory, computational geometry (space filling diagrams), and computational topology (filtrations, persistence diagrams, Betti numbers). Readers not familiar with these fields are referred to SI section 8.

### 2.1 Structural motifs

Let A and B be the two structures to be compared. Without loss of generality, we assume that the particles considered are *C_α_* atoms, denoted *A* = {*a_i_*} and *B* = {*b_i_*}. We define:

#### Definition. 1

*A* motif *shared by A and B is a pair of set of particles M^(A)^ ⊂ A and M^(B)^ ⊂ S_B_ of the same size, together with an alignment, that is a one-to-one correspondence between their a.a.*.

The alignment allows computing the least RMSD, lRMSD(*M_A_, M_B_*). We further compare this quantity to that of the structures themselves:

#### Definition. 2

*(least RMSD ratio) Consider two structures A and B, and assume that a structural alignment between them has been computed, together with the corresponding lRMSD(A, B). The* least RMSD ratio *of the motif is defined by:*

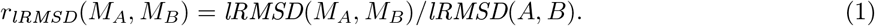

*The sets M_A_ and M_B_ are called* structural motifs *provided that* |*M_A_*| = |*M_B_*| ≥ *τ_MS_ and r_lRMSD_*(*M_A_, M_B_*) ≤ *τ/, for appropriate thresholds τ_MS_* (> 0) *and τ*/(∈ (0,1)).

**Remark 1** *We note that the alignment used in Def. 1 depends on the context-homologous proteins versus conformations. In the former case, we use two methods respectively favoring long and flexible alignments (`Kpax`, [12], and backbone geometry (`Apurva`, [17]). In the latter case, the alignment is the trivial identity alignment. That is, we retain the a.a. common to both conformations–as some may be missing e.g. loops in crystal structures*.

**Remark 2** *Motifs are local structural patterns. They can be combined to compare whole structures using the notion of combined RMSD_comb._, as recalled in SI Section 8 and [22]*.

### 2.2 Structural motifs from alignments: overview

Let us first convey the intuition of our method to compare two structures *A* and *B*. For the time being, consider the simple case where the two structures correspond to two conformations of a molecule consisting of say two domains connected by a flexible linker. If the relative position of the two domains changes in switching from one conformation to the other, the only conserved distances are those involving any two *C_α_* from the same domain. Assume that we have computed a score assessing the conservation of the distance between two *C_α_*. A natural strategy to identify the two domains therefore consists in incrementally processing scores, and maintaining the connected components linking the a.a. involved in the scores, using a union-find data structure [25]. We implement this idea via a four step method (Fig. 1):

**Step 1: Computing the seed alignment and its scores.** We first compute a *seed* structural alignment between the two structures *A* and *B*. If the two structures are two conformations of the same molecules, the alignment is trivially the identity. We then compute the distance difference matrix associated with the alignment, so as to qualify the conservation of distances between *C_α_* between the two structures.
**Step 2: Building the filtration and its persistence diagram.** We build the conserved distances and space filling diagram filtrations already mentioned, together with the associated persistence diagrams, denoted PD_*A*_ and PD_*B*_ for the two molecules. Points on these PD correspond to regions in the structures.
**Step 3: Computing structural motifs.** We identify relevant pairs (*a, b*) of points, with *a* ∈ PD_*A*_ (resp. *b* ∈ PD_*B*_). We compute a structural alignment involving the connected components of the sublevels sets associated with these points, from which we compute a structural alignment and retrieve structural motifs complying with Eq. (1).
**Step 4: Filtering structural motifs.** Motifs are filtered to get rid of redundancy (inclusion), and to retain those which are statistically significant.

**Figure 1:**
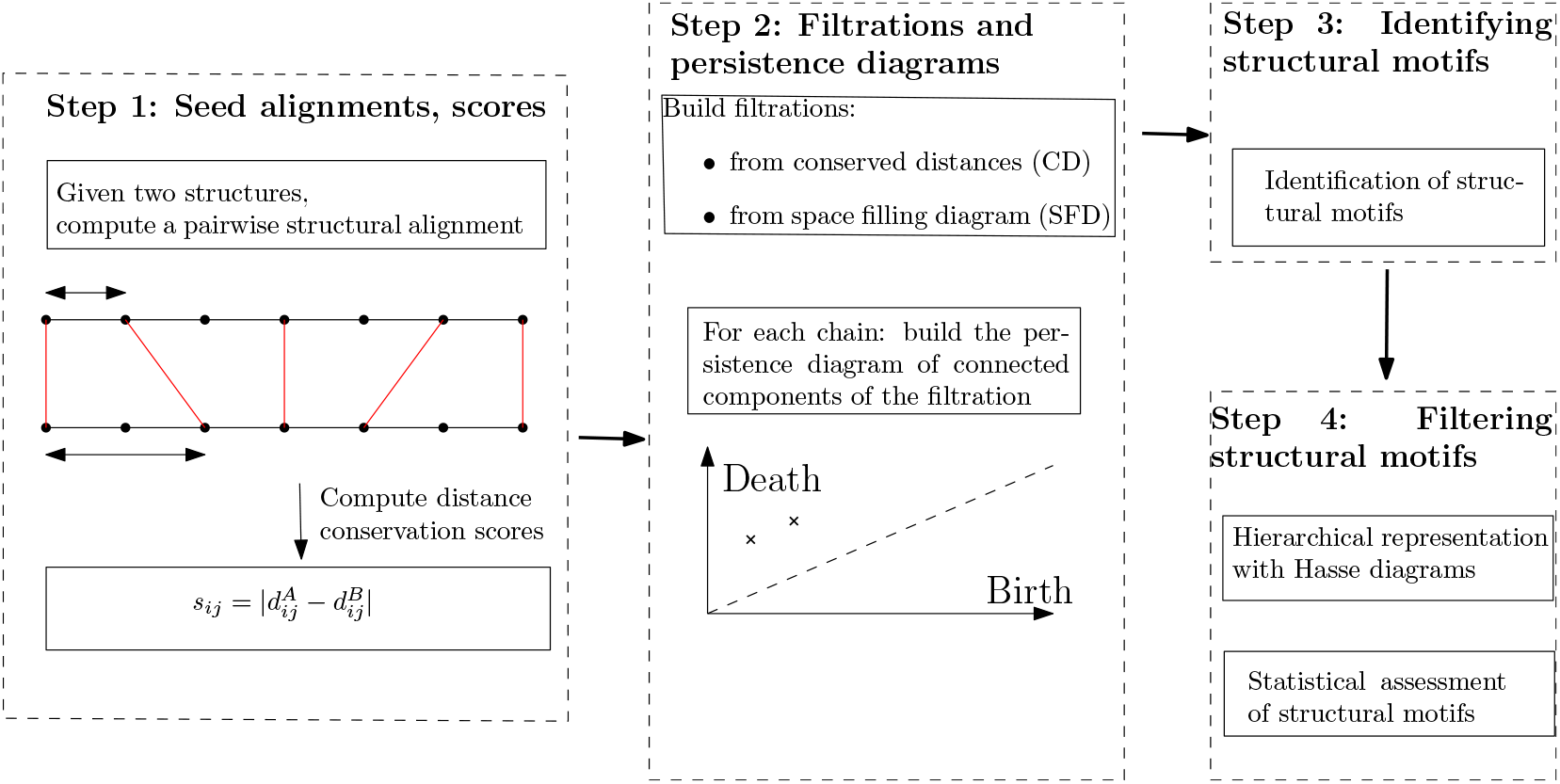
The four step method to identify structural motifs

We now detail these steps.

### 2.3 Step 1: Computing the seed alignment and its scores

Consider a structural alignment between the two structures *A* and *B*. We denote 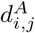 the distance between the *C_α_* carbons of indices *i* and *j* on chain *A*, and likewise on chain *B*. These quantities are used to define the *distance difference matrix (DDM)*, a *N* × *N* symmetric matrix whose entries, also called *scores*, are given by:

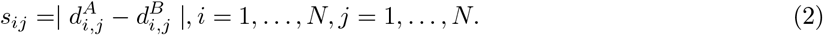

### 2.4 Step 2: Building the filtration and its persistence diagram

We use the 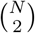 scores of Eq. (2) to define two filtrations. The first one exploits the ordering of a.a. pairs induced by scores, to perform a multiscale analysis of structurally conserved pairs of a.a. The second one exploits an ordering of individual a.a. induced by scores, to also perform a multiscale analysis of structurally conserved regions.

#### 2.4.1 Conserved distances filtration (CD filtration)

##### Score index

Consider the sorted sequences of scores from Eq. (2). An edge involves two a.a., so that in processing edges by increasing scores, we iteratively connect a.a. This processing is actually Kruskal’s algorithm to build a minimum spanning tree (MST) [26]. We define:

###### Definition. 3

*(Score index) The scores contributing to a MST connecting the a.a. are called* non redundant. *Denoting N the alignment size, there are N − 1 such scores. The* score index *of a non-redundant score is its index in the sequence of non redundant scores*.

##### Filtration

Our rationale is to identify rigid domains by incrementally processing scores contributing to the MST. Recall that Kruskal’s algorithm processes edges by increasing scores. The connexion between connected components of the MST as characterized by the first and second Betti numbers *β*_0_ and *β*_1_, and scores/edges as processed in Kruskal’s algorithm is as follows:

- (c.c. creation) edge triggering the creation of a connected component: the score involves two *C_α_* not processed previously; this score creates a new c.c. in the MST. In terms of topology of the MST: *β*_0_+ = 1.
- (accretion) edge triggering accretion: the score involves only one *C_α_* not processed previously, which therefore contributes to the extension of an existing c.c. In terms of topology, neither *β*_0_ nor *β*_1_ change.
- (c.c. destruction) edge triggering the destruction of a c.c.: the score involves two *C_α_* already involved in scores processed previously, and these *C_α_* belong to two different c.c.. These c.c. merge, so that *β*_0_− = 1. (NB: if the two *C_α_* belong to the same c.c., then *β*_0_ remains unchanged but *β*_1_+ = 1. Such edges creating cycles and are not used in the MST.)

##### Persistence diagram

The persistence diagram codes the lifetime of c.c. in the MST during its construction. Note that the persistence diagram constructed is the same for the two structures.

**Remark 3** *The number of points in the PD is equal to the alignment length N. To see why, note that in running Kruskal’s algorithm, vertices are associated with two types of* creation *events. For an edge triggering a c.c. creation: two c.c. are created-one for each vertex, but one dies immediately and has a null persistence. For an edge triggering accretion, one c.c. corresponding to the new vertex is created, and this c.c. also has a null persistence. Finally: every vertex of the graph is involved in exactly one of these two edge types*.

#### 2.4.2 Space filling diagram filtration (SFD filtration)

We now exploit the scores *s_ij_* in a different way, by focusing on individual *C_α_* rather than edges. We assign assign to each *C_α_* carbon an integer in the range 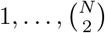, called its *rank*.

*C_α_* **ranks.** We assess the structural conservation of the neighborhood of a given *C_α_* with (Fig. 2):

#### Definition. 4

*(C_α_ rank) The* absolute *C_α_* rank *of a C_α_ is the smallest index 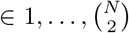 of the score s_ij_ involving this C_α_. The largest absolute C_α_ rank is denoted N_r_, and the number of (distinct) absolute C_α_ ranks n_r_*.

*The C_α_* rank *of a C_α_ is the index of its absolute C_α_ rank - an integer in the range 1,…, n_r_*.

*The amino-acids of structures A and B whose C_α_ ranks are equal to i are denoted A_(i)_ and B_(i)_, respectively*.

**Figure 2:**
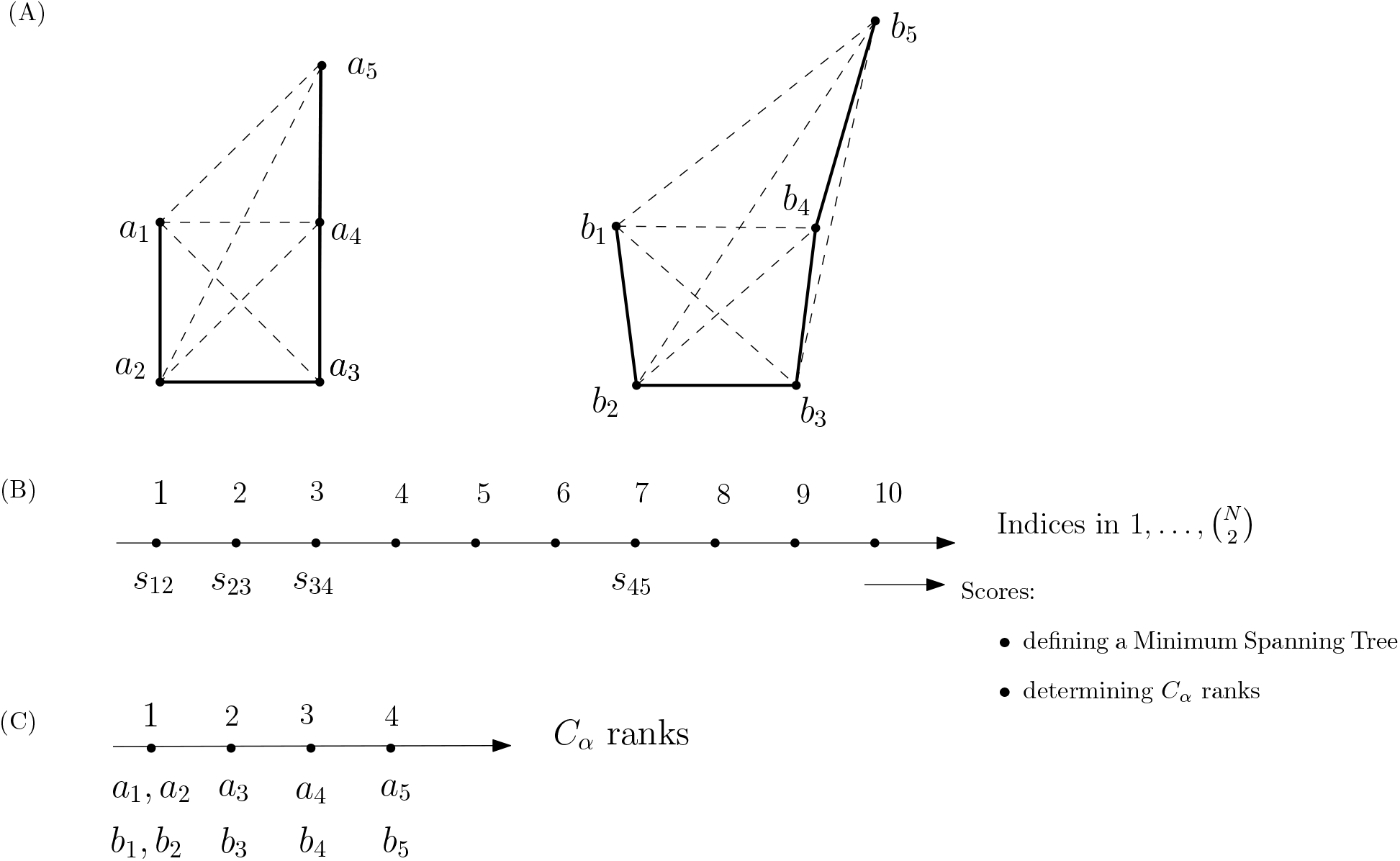
Step one, Computing the seed alignment and its scores: method. (**A**) Consider the alignment (*a_i_* ↔ *b_i_*)_*i* = 1, …, _*N*__ between two fictitious chains (bold line-segments) of length *N*= 5. (**B**) The 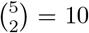 scores are sorted. The scores involved in the definition of the conserved distances (CD) filtration, which also define a spanning tree connecting the a.a. of each structure, are: *s*_1,2_, *s*_2,3_, *s*_3,4_, *s*_4,5_. (**C**) On this toy example, the same scores contribute to the definition of *C_α_* ranks, from which the space filling diagram filtration is defined.

Note that absolute *C_α_* ranks are identical for atoms mapped with one another. We summarize with the following:

**Observation. 1** *The following upper and lower bounds are tight*:

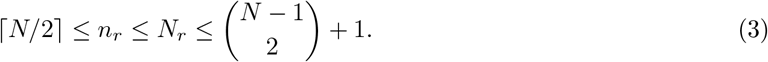

##### Proof.

Consider first absolute *C_α_* ranks. For the lower bound, since a given score contributes to at most two ranks, 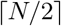 scores are needed to cover the *N C_α_*.

For the upper bound, consider a (fictitious) molecule with a rigid region of *N* − 1 *C_α_−s*, and one mobile *C_α_.* The absolute *C_α_* rank of the mobile atom is equal to 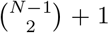 since it is determined by the most conserved distance out of the N — 1 distances to the *C_α_−s* of the rigid region. This number is worst-case, since for the *C_α_* with largest *C_α_* rank, out of the *N* − 1 distances to the remaining *C_α_−s*, only the smallest one determines its *C_α_* rank.

The bounds *C_α_* ranks are trivial given those on absolute *C_α_* ranks. □

##### Filtration

Consider now the processing of a.a. by increasing *C_α_* rank. We maintain for each molecule a SFD, and report contacts between a.a. using the associated α-complex (section 8.3). Consider a SFD (that of molecule *A* or *B*). A *C_α_* may be ascribed to the following categories, which we distinguish in terms of topological changes:

- (c.c. creation) The *C_α_* creates a new c.c. of the SFD; that is, *β*_0_ + = 1.
- (c.c. destruction) The *C_α_* bridges the gap between two existing c.c.; that is, *β_0_− =* 1.
- (cycle creation) The *C_α_* creates a cycle in an existing c.c.; that is, *β_1_+ =* 1.
- (accretion) The *C_α_* contributes to the enlargement of an existing c.c.; neither *β*_0_ nor *β*_1_ change.

##### Persistence diagram

Consider chain *A* - we proceed mutatis mutandis for chain *B*. Using the ordering of a.a. defined by *C_α_* ranks, let 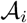 be the set of a.a. whose rank is at most *i*, that is 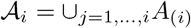. We denote the SFD of this collection of balls by 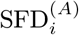. The connected components of this SFD evolve upon inserting a.a. (SI Fig. 12). Because the whole polypeptide chain defines a connected domain, each c.c. but the one created by *A(_1_)* is characterized by two dates, namely its birth date and it death date. Furthermore:

**Observation. 2** *Consider two chains A and B whose space filling diagrams are connected. Upon processing the last C_α_ rank, the space filling models 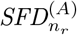 are 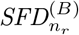 are connected*.

Note, however, that as opposed to the PD obtained with scores, those obtained with *C_α_* ranks are not identical. Indeed, the connectivity obtained along the filtration depends on the relative position of a.a., which differ in *A* and *B*.

#### 2.4.3 Comments

##### Accretion

The two types of filtrations undergo different topological events, as characterized by the evolution of Betti numbers. One common feature though is the presence of *accretion*: accretion is the addition of one edge (conserved distances filtration) or one a.a. (SFD filtration), without any change in *β*_0_ or *β*_1_.

Accretion corresponds to the incremental formation of a motif, in such a way that no motif is *stable*. In topological persistence terms, accretion is characterized by null persistence.

Practically, accretion calls for two comments:

- While accretion does occur, our experiments show that its extent is not such that motifs with significant persistence do not exist.
- As we shall see, iterative alignments provide a natural way to rescue regions of motifs plagued by accretion.

##### Plots

To study the correlation between *C_α_* ranks and properties of the sequences, we define (Fig. 3):

###### Definition. 5

- Score plot: *score s_ij_ as a function of the C_α_ rank.*
- Sequence shift plot: *for chain A (or chain B), the function j − i (distance along the sequence) as a function of the C_α_ rank.*
- *C_α_* distance plot: *for chain A, the function 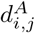 (or 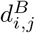) as a function of the C_α_ rank*.

**Figure 3:**
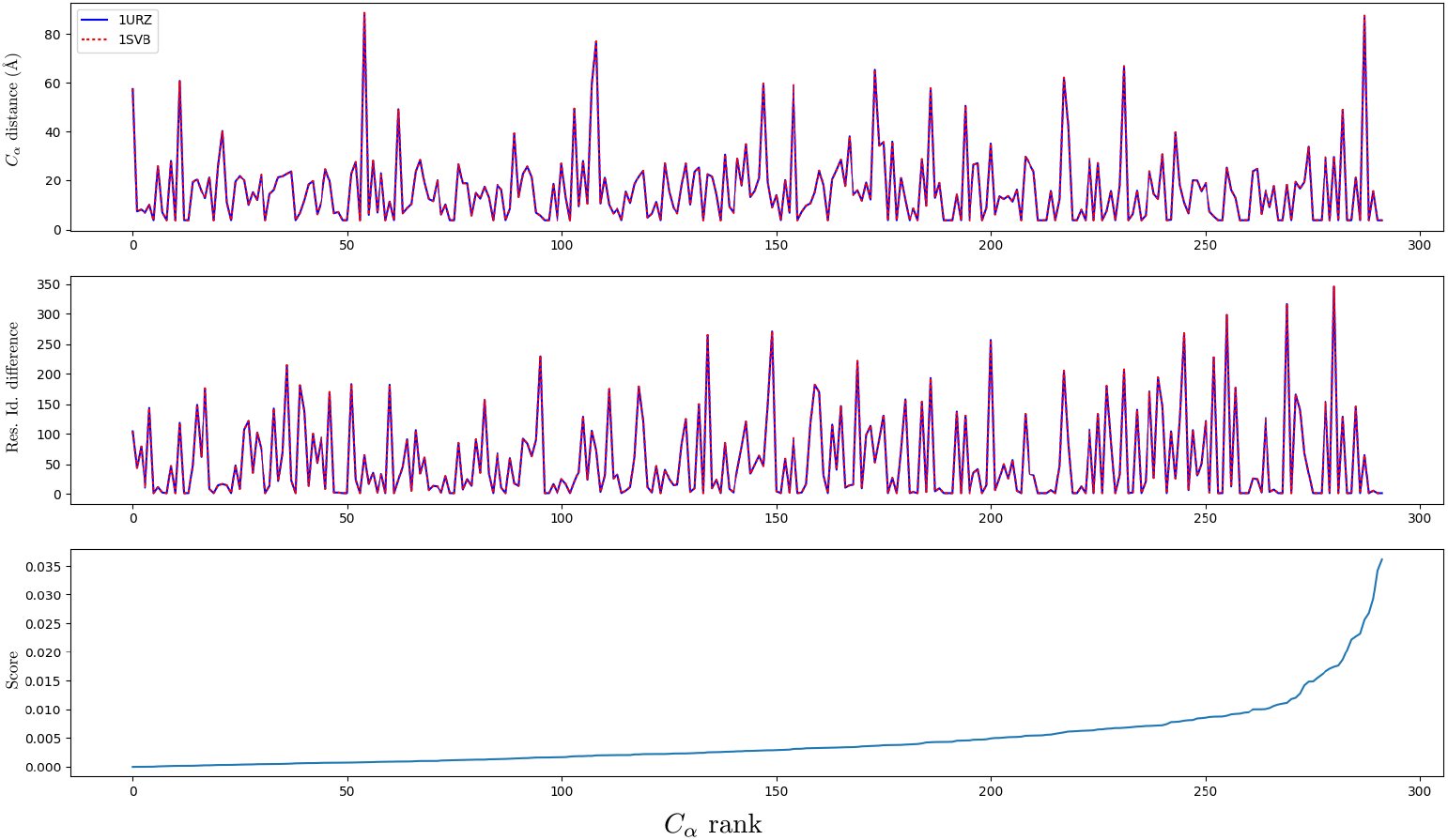
Step 1, Computing the seed alignment and its scores: illustration for a class II fusion protein of TBEV in two different conformations (pre-fusion (PDB: 1SVB), post-fusion (PDB: 1URZ)) Plots definition: see Def. 5; overview of structures, see SI Fig. 11. (**Top**) *C_α_* distance plot (**Middle**) Sequence shift plot (**Bottom**) Score plot No correlation is observed between *C_α_* ranks and (i) the proximity along the sequence, and (ii) the location on SSE. That is, *C_α_* ranks identify rigid pairs throughout the structures.

### 2.5 Step 3: Computing structural motifs

This step exploits either of the aforementioned persistence diagrams. It also relies on an alignment algorithm, which is assumed to be identical to the one used to compute the seed alignment and the scores.

#### From connected components to motifs

Consider a point *a* = (*b_a_, d_a_*) ∈ PD_*A*_, with *b_a_* and *d_a_* the birth and death dates of *a*, respectively. To point *a*, we associate the set of connected component of the sublevel set defined by *d_a_*, which we denote *F_A_*(*a*) = {*c*_1_, …, *c_n_A__*}. For *b* ∈ PD_*B*_, we obtain similarly the set 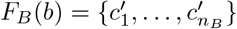.

##### Definition. 6

*(Structural motif) Consider two points from the PD, namely a ∈ PD_A_ and b ∈ PD_B_, and the associated connected components of sublevel sets, i.e. F_A_(a) = {c_1_, …, c_n_A__} of* 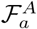 *and* 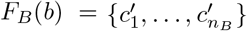.

*Consider pair of large enough c.c., that is* 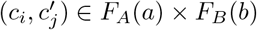 *with* 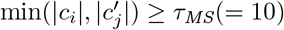.

*A* structural motif *for that pair is defined by the structural alignment obtained for* 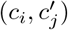.

It should be noticed that since a motif consists of a.a. singled out by a local alignment, it may not be connected in 3D space, even when the persistence diagram is associated with the SFD filtration.

#### Localizing comparisons

Definition 6 requires processing all pairs of c.c. in *F_A_*(*a*) × *F_B_(b)*. On the other hand, under suitable assumptions, persistence diagrams are known to be stable [27]. As a heuristic to reduce the number of points from PD_*A*_ and PD_*B*_ compared, we require the Euclidean distance in PD space is less than a threshold *τ*:

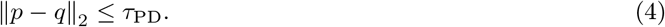

#### The case of homologous proteins

In comparing homologous proteins, to reduce the number of pairs of c.c. processed, we further require the two compared c.c. to have comparable size, that is:

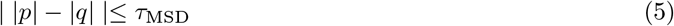

#### The case of two conformations

When comparing two conformations of the same structure, the situation is easier since one has the identical alignment between chains of the two conformations, see Remark 1. In that case, we perform a one step clustering of c.c. using the D-family matching algorithm [28] - a method to group clusters into meta-clusters with a one-to-one correspondence between meta-clusters. We then process the pairs corresponding to meta-clusters.

### 2.6 Step 4: Filtering structural motifs

#### Motif inclusion

Consider the subsequences of a.a. defining motifs. Due to the nested structure of sets obtained from filtrations, motifs may be nested. Since a motif reduces to a set of a.a., we may consider the partial order defined by set inclusion, and the associated Hasse diagram. (NB: the Hasse diagram is the directed graph whose edges precisely code inclusion between motifs.) Of particular interest are the terminal nodes of the Hasse diagram, which may be seen as maximal motifs in terms of size.

#### Statistical significance

The statistical significance of our motifs can also be assessed using a two-sample test involving random motifs (SI Section 9).

### 2.7 Expanding motifs with iterative aligners

Iterative alignments [14, 12] consist in iteratively finding the alignment for fixed positions of the conformations-via dynamic programming, and finding optimal superimposition given the alignment–the classical rigid superimposition problem. The process is typically iterated until a fixed point or a cycle is reached. Seeding such alignments with our motifs is especially interesting for two reasons. First, the construction of motifs inherently copes with accretion–Sec. 2.4.3. Therefore, iterative alignments allow the extension of motifs with a.a. involved in accretion. Second, motifs define motif graphs (Def. 7, SI Sect. 8.1). Given a c.c. of a motif graph, an iterative alignment can be seeded with each motif from this cc. In the end, we retain the alignment maximizing the G-score [12].

It should be noted that when processing two conformations, the trivial/identity alignment is used (remark 1), and the process simplifies to the following two steps:

1. (Alignment given relative position) Add to the current alignment each residue pair whose distance is below a threshold *τ_L_*.
2. (Relative position given alignment) Use the previously computed alignment to perform a rigid registration.

## 3 Implementation

All methods are available in the Structural Bioinformatics Library (http://sbl.inria.fr, [29]).

### 3.1 Seed aligners

A structural alignment algorithm is required to get the seed alignment and to compare two points from the persistence diagrams. Practically, we use the aforementioned state-of-the-art options `Apurva` [17] (`SBL` package: https://sbl.inria.fr/doc/Apurva-user-manual.html) and `Kpax` [12] (SBL package: https://sbl.inria.fr/doc/Iterative_alignment-user-manual.html).

### 3.2 Persistence diagrams

Consider the score filtration—Sec. 2.4.1. We build a complete edge weighted graph whose nodes are the a.a.. The weight of an edge is the score associated to this edge Eq. (2). Sorting the edges of this graph yields a filtration, from which the persistence diagram for connected components is easily maintained using a union-find data structure. Practically, we use the Morse theory based analyzer from the `SBL` (`SBL` package: https://sbl.inria.fr/doc/group__Morse__theory__based__analyzer-package.html).

Consider the SFD filtration-2.4.2. We build a vertex weighted graph whose nodes are the a.a., sorted by increasing *C_α_* rank. For each molecule, we sequentially insert the a.a., maintain the corresponding *α*-complex, and build the associated persistence diagram, also using the Morse theory based analyzer from the `SBL`.

### 3.3 Motifs

#### Methods: Non-iterative aligners

Combining the two seed aligners (`Apurva, Kpax`) and the two persistence diagrams (CD: conserved distances, SFD: space filling diagram) yields four options. One also needs to distinguish conformations versus homologous proteins, as the former involve the trivial identity alignment (remark 1). Summarizing, we report in Experiments results for the following combinations (in parenthesis, the color used for plots when appropriate):

- Aligners requiring an alignment: `Align-Kpax-CD, Align-Kpax-SFD, Align-Apurva-CD, Align-Apurva-SFD;`
- Aligners with the identity alignment: `Align-Identity-CD, Align-Identity-SFD`.

Practically, these methods are implemented within the following executables from the SBL-each giving access to the CD and SFD filtrations: `sbl-structural-motifs-chains-apurva.exe, sbl-structural-motifs-chains-kpax.exe, sbl-structural-motifs-conformations.exe`.

#### Methods: Iterative aligners

Seeding `Kpax` with our motif yields four different iterative aligners, namely `Align-Kpax-SFD`/iter, `Align-Kpax-CD`/iter, `Align-Apurva-SFD`/iter, `Align-Apurva-CD`/iter.

#### Packages from the SBL and main parameters

Methods and executables are summarized in SI Table 2. The programs computing motifs are provided in the package Structural_motifs package (`SBL` package: https://sbl.inria.fr/doc/Structural_motifs-user-manual.html), and are used in particular to compute flexible distance measures in the package Molecular_distances_flexible (https://sbl.inria.fr/doc/Molecular_distances_flexible-user-manual.html). Iterative aligners are provided in the package Iter-ative_alignment (https://sbl.inria.fr/doc/Iterative_alignment-user-manual.html).

The main parameters of these programs are: *τ*_MS_ for the motif size (Def. 6), *τ*/ for the lRMSD ratio (Def. 2), *τ*_PD_ for the comparison of persistence diagrams (Eq. 4). Practically, the following parameters are used:

- comparing homologous proteins: *τ*_MS_ = 10, *τ*/ = 0.8, *τ*_MSD_ = 10; SFD filtration: *τ*_PD_ = 20; CD filtration: *τ*_PD_= 0.
- comparing conformations: *τ*_MS_ = 10, *τ*/ = 0.5; SFD filtration: *τ*_PD_ = 5; CD filtration: *τ*_PD_ = 0.

## 4 Results

### 4.1 Datasets

#### Structural comparisons and motifs

To understand properties of motifs, we use a dataset of eight class II fusion proteins (SI Fig. 11), namely proteins used by enveloped viruses to trigger the fusion between their membrane and that of the target cell [30]. These 8 structures (sizes varying from 380 to 461 a.a., SI Fig. 11) yield 8 × 7/2 = 28 pairwise comparisons. The rationale for this dataset is twofold. Structurally, class II fusion proteins are elongated molecules with three domains (DI, DII, DIII) composed primarily of *β*-sheets. The central DI domain connects via a flexible hinge to the longer DII. Typically, DII contains several conserved disulfide bonds as well as the so-called fusion loop at its tip. Additionally, a linker region connects DI to the DIII domain, which has an Immunoglobulin (Ig)-like fold.

On the biological side, (class II) fusion proteins are challenging structures, as they harbor a low sequence identity (typically less than 20%), and overall loose structural homology (typically of the order of 10Å). Yet, they have the same function, for a reason which has remained elusive. Thus, in investigating structurally conserved motifs of such proteins, we provide insights on fusion mechanisms [31].

On the computational side, such a dataset remains tractable. To see why, recall that we use two aligners, namely Apurva (with a focus on contacts) and Kpax (with a focus on backbone geometry), at two stages: first, to obtain the seed alignment; second, to obtain the motifs by performing a local alignment when processing pairs of points from the persistence diagrams (Def. 6). This latter operation is the computational bottleneck. While Kpax is fast and typically runs under a minute even on large examples, Apurva may be quite slow even on small examples. With default parameters, run time can reach one hour on a single instance. The comparison of two PD may thus take several hours, which greatly limits the size of the data set when performing large scale comparisons.

#### Comparison against flexible aligners

For a direct comparison against flexible aligners, we use the 10 ‘difficult’ structures from [23].

### 4.2 Method illustration: investigating a conformational change

We illustrate each step of our method to study the conformational change undergone a prototypical class II fusion protein, from the tick-borne encephalitis virus. The ectodomain of this protein was crystallized both in soluble form (PDB: 1SVB, [32], 395 residues) and in postfusion conformation (PDB: 1URZ, [33], 400 residues). For these structures, the identity alignment identifies 376 a.a.

**Step 1 – scores and *C_α_* ranks.** We compute *C_α_* ranks as described in Sec. 2.3 and analyze the *C_α_*distance plot, the sequence shift plot, and the score plot. (Fig. 3). No correlation is observed between *C_α_*ranks and (i) the proximity along the sequence, and (ii) the location on SSE (inferred from *C_α_* distances). That is, *C_α_* ranks identify rigid pairs throughout the structures.

**Step 2 – persistence diagrams.** Persistence diagrams exhibit different amounts of persistent events / accretion (SFD filtration, Fig. 4; CD filtration, SI Fig. 13). When using a SFD filtration, most main merging events happen before *C_α_* rank 100. As expected, in the case of conserved distances, the PD is denser and less prone to accretion.

**Figure 4:**
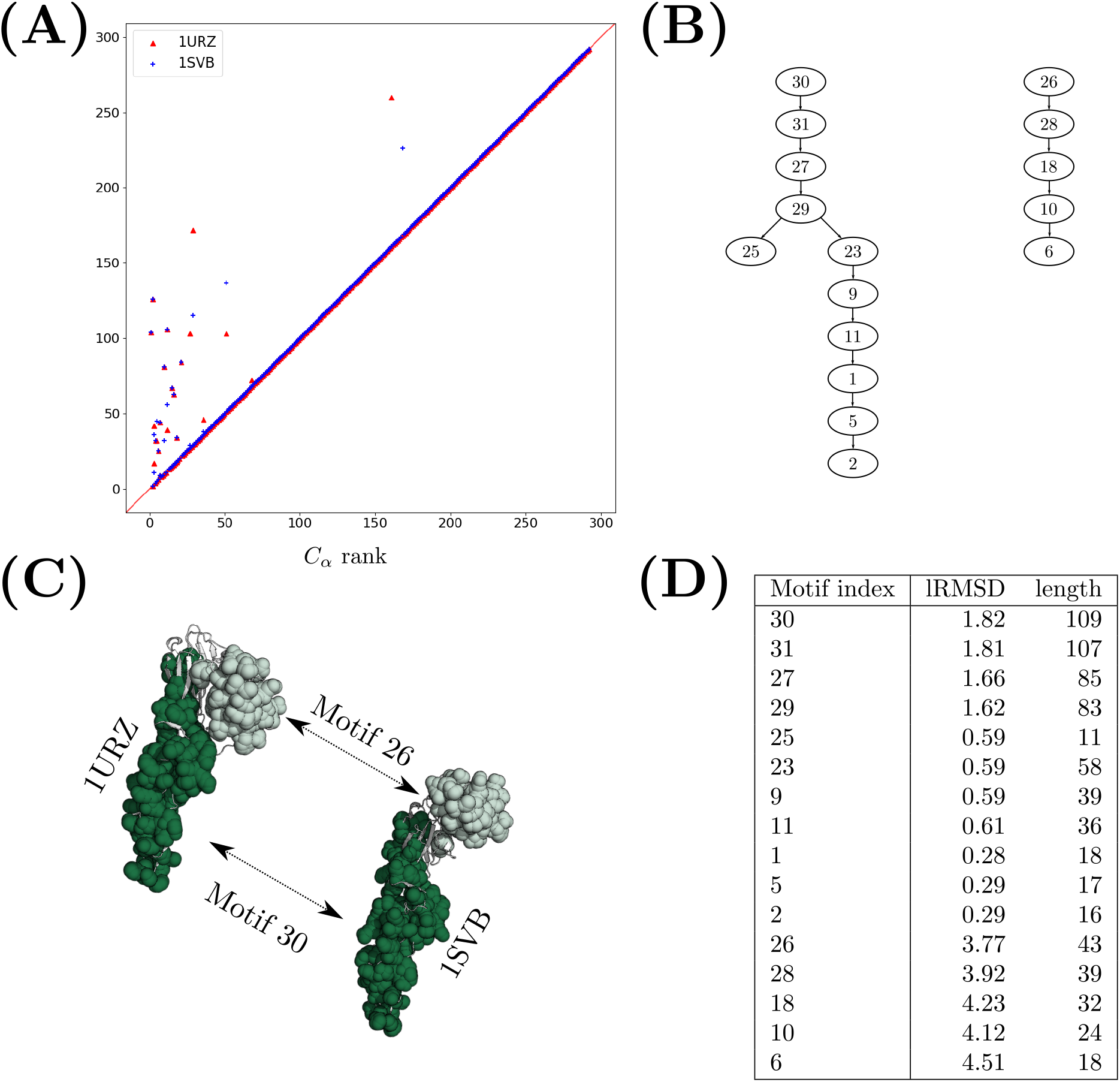
Step 2, Building the filtration and its persistence diagram: illustration for 1URZ-1SVB with Align-Identity-SFD. Comparing two conformations of TBEV class II fusion protein yields two nested sets of structural motifs which successfully characterize the two parts of the hinge motion. (**A**) Persistence diagram for SFD filtration. (**B**) Hasse diagram of structural motifs. Each motif has its unique index. (**C**) Selected motifs. Each motif corresponds to one part of the hinge motion associated to the two conformations. (**D**) Statistics for the structural motifs.

**Step 3 and 4 – motifs.** The hierarchical structure of the motifs found is coded in the Hasse diagram coding motif inclusion (Sec. 2.6; Fig. 4). On this example, the two motifs at the root of the Hasse diagram characterize the hinge motion associated with the conformational change (Fig. 4(c)). This hinge motion is best characterized by comparing the global lRMSD yielded by our seed aligners, and the combined RMSD which mixes the lRMSD of the two motifs (Eq. 6, [22]):

- `Kpax`: #a.a.: 296, lRMSD : 9.28 Å;
- `Apurva`: #a.a.: 370; lRMSD : 11.1 Å;
- Motifs yielded by `Align-Identity-SFD`: #a.a.: 152; RMSD_Comb._ : 2.53 Å.
- Motifs yielded by `Align-Identity-CD`: #a.a.: 161; RMSD_Comb._ : 1.26 Å.

#### Iterative alignment

Motifs can be used to seed iterative aligners-Sec. 2.7. Using the SFD filtration with default parameters, `Align-Identity-SFD`/iter (See SI Table 2) identifies the rigid groups formed by the DI and DII domains on the one hand, and the DIII domain on the other. These motifs are characterized by (Fig. 4):

- Motif 1 (DI, DII): #a.a.: 171, lRMSD : 0.86.
- Motif 2 (DIII): #a.a.: 93, lRMSD : 0.58
- Overall comparison: #a.a.: 264, RMSD_Comb_. : 0.76

Using the CD filtration, `Align-Identity-CD`/iter with tuned parameters *τ*_PD_ = 0, *τ*_MS_ = 5, *τ*/ = 0.5, *τ_L_* = 0.8 (Nb: *τ_L_* specific to iterative aligners for conformations). even identifies all three domains of the structure (Fig. 5). These motifs are characterized by:

- Motif 1 (DII): #a.a.: 115, lRMSD : 0.51
- Motif 2 (DI): #a.a.: 79, lRMSD : 0.37
- Motif 3 (DIII): #a.a.: 40, lRMSD : 0.49
- Overall comparison: #a.a.: 234, RMSD_Comb_. : 0.46

**Figure 5:**
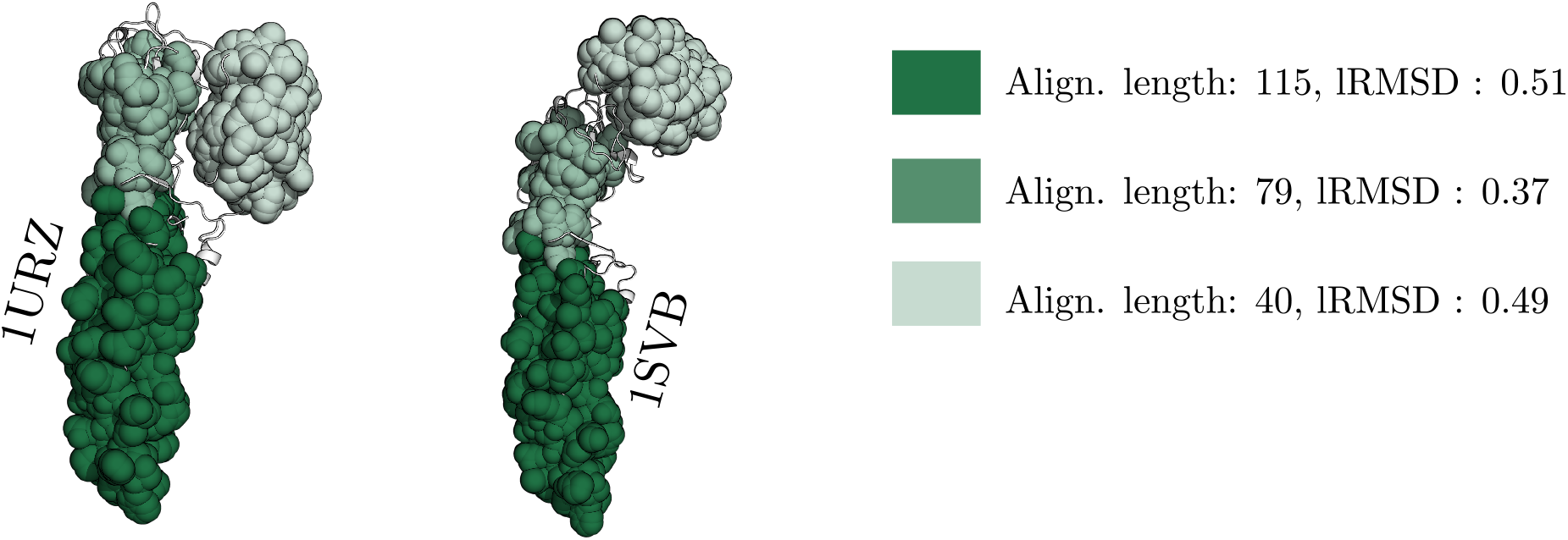
Using motifs found with `Align-Identity-CD` as seeds for an iterative alignment characterizes all three domains of the TBEV class II fusion proteins.

As illustrated by this example, using motifs in combination with the combined RMSD yields a reduction of ~ 20 of the metric used to perform the structural comparison.

### 4.3 Motifs: a case study for homologous proteins

#### 4.3.1 Comparing the four aligners: overall comparison

##### Method

We challenge our four methods (`Align-Kpax-CD, Align-Kpax-SFD, Align-Apurva-CD, Align-Apurva-SFD`) and also compare them to the seed aligners (`Apurva, Kpax`), using all pairwise comparisons of the class II fusion dataset. In a first step, we analyze statistics describing the motifs yielded by the contenders. In a second step, we exploit motifs to perform an overall comparison of whole structures based on motifs. For the seed aligners `Apurva` and `Kpax`, we report the lRMSD associated with the seed alignment, together with the alignment length. For our four aligners, we report the combined RMSD_Comb_. associated with our motifs and the number of a.a. involved in this calculation [22].

##### Seed aligners

The alignments returned by our base aligners `Apurva` and `Kpax` for the 28 comparisons call for two comments (Fig. 6). First, `Apurva` alignments have a lRMSD which can be up to four times larger than `Kpax` alignments; second, `Kpax` alignments are smaller -the median is ≃ 100 residues smaller. This is in line with the goals of these aligners, as `Apurva` tries to maximize the numbers of contacts, while `Kpax` favors the coherence of backbone geometry.

**Figure 6:**
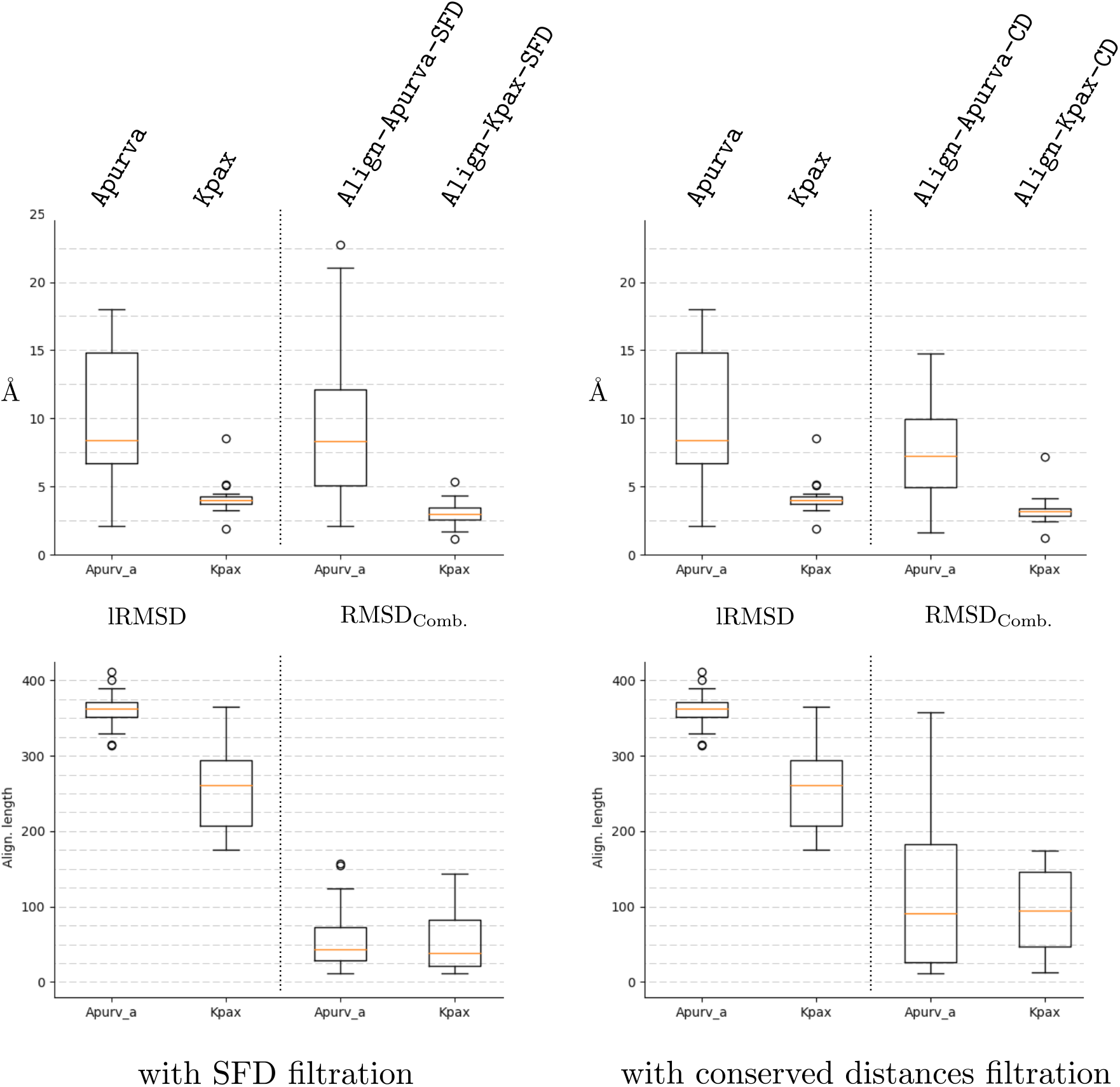
Motif based comparison of the four aligners Align-Kpax-CD, Align-Kpax-SFD, Align-Apurva-CD, Align-Apurva-SFD. Note that an aligner is defined by the conjunction of an alignment method (`Apurva, Kpax`) and a filtration method (SFD: Space Filling Diagram; CD: conserved distances). The comparison is based on two statistics: for seed aligners (`Apurva, Kpax`), the lRMSD of the alignment and the alignment size; four our four aligners: the combined RMSD RMSD_Comb_. defined from the motif graph, and the number of a.a. involved (the number of vertices of the motif graph).

##### Motifs per se

The motifs returned by our four methods provide insights on the merits of the seed aligners and the filtrations (see also SI Sec. 10.1 for the statistical significance of motifs.) Three facts emerge from the general statistics obtained on the whole dataset (Table 1).

**Table 1:**
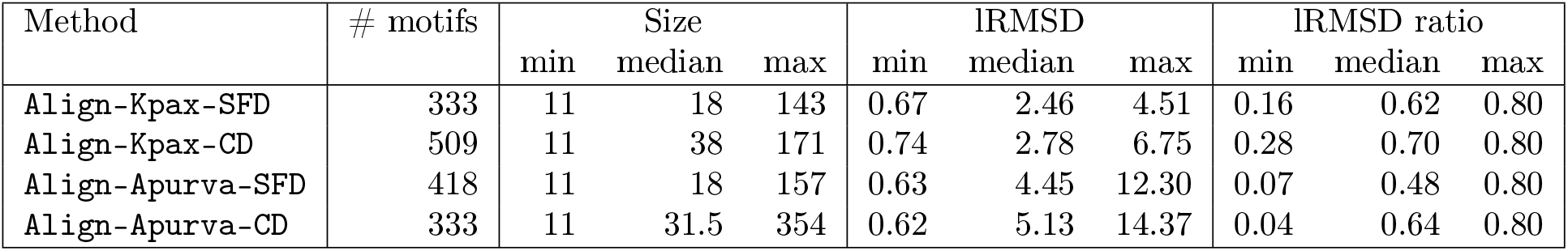
General statistics for motifs returned by our four methods. Statistics are reported for the 28 pairwise comparisons on the class II fusion proteins.

First, the CD filtration yields longer motifs (Table 1; median size 38 and 31.5, versus 18 and 18). This is expected, as defining connected components from a space filling diagram is more stringent than from the MST exploiting conserved distances. Second, the lRMSD of motifs returned by methods using `Apurva` as seed aligner are larger (Table 1; median values of 4.45 and 5.13, against 2.46 and 2.78). This is also expected, as the seed alignment of `Apurva` favors length, while that of `Kpax` favor geometric coherence. The third is that the lRMSD ratio of motifs returned by methods using `Apurva` tends to be smaller (Table 1; median values of 0.48 and 0.64, against .62 and 0.70). This stresses the importance of the seed alignment size in the definition of the lRMSD ratio (Def. 2).

##### Motifs to compare whole structures

The lRMSD of motifs can be combined to compare whole structures [22] (Fig. 6). The RMSD_Comb_. is always globally better than the lRMSD, except in the case of `Align-Apurva-SFD` in which it performs poorly in few instances (Fig. 6, top left). For `Align-Kpax-SFD` and `Align-Kpax-CD`, the median RMSD_Comb_. is ≃ 1 Å smaller than the initial lRMSD median. For `Align-Apurva-SFD`, the median RMSD_Comb_. is comparable to the initial lRMSD median but both the top and lower quartile are ≃ 2.5 smaller so that the results are more homogeneous. When using a SFD filtration, the median number of residues involved in the RMSD_Comb_. is ≃ 50 residues. In some instances, it can be an order of magnitude (between 30 and 40 times) smaller than the seed alignment length (Fig. 6, bottom left). With conserved distances, the median number of residues involved in RMSD_Comb_. is ≃ 100 residues.

#### 4.3.2 Comparing the four aligners: pairwise comparison

##### Method

We now compare the four methods, exploiting the multiscale nature of the motifs returned. Each motif is characterized by a 2D point (motif size, lRMSD). The comparison of two structures can therefore by summarized in this space by a point cloud and the associated Pareto envelope. Note that in (motif size, lRMSD) space, the Pareto front is defined using the following notion of domination: a motif *a* dominates a motif *b* iff *a* has more a.a. than *b*, and the lRMSD of *a* is smaller than that of *b*.

In the sequel, we compare the Pareto fronts of the four methods for each of the 28 pairwise comparisons, in order to assess whether a particular method stands out (for the full matrix of Pareto fronts, see SI Fig. 10.1).

Each point cloud and curve is color coded with respect to one of the four methods: `Align-Kpax-CD` (blue), `Align-Kpax-SFD` (orange), `Align-Apurva-CD` (green), `Align-Apurva-SFD` (red).

##### Results

Four scenarios emerge from the 28 pairwise comparisons.

1. **Blue dominates (Fig. 7):** this is the most common case. Generally, `Align-Kpax-CD` performs better than other methods w.r.t lRMSD and motif sizes. In the example, even though the blue motif is ≃ 2× larger than the orange motif (which is itself significantly larger than any other motif), its lRMSD is slightly lower.
2. **Mixed domination, green then blue (Fig. 8):** About one third of the comparisons show a relayed domination between a given method and `Align-Apurva-CD` (green). The example from Fig. 8 is particularly striking as `Align-Apurva-CD` returns very large motifs (343 residues).
3. **Orange dominates (Fig. 9):** In a few other cases, `Align-Kpax-SFD` (orange) dominates. The example from Fig. 9 shows the particularities of each method. The blue motif (found by `Align-Kpax-CD`) is composed of many smaller connected components distributed through out the structure. On the other hand, the orange motif (found by `Align-Kpax-SFD`) is localized on the top of the structure and formed of a unique, larger, connected component.
4. **Mixed domination, all (Fig. 10):** Finally, in some cases, we observed a four-way relayed domination in which each method dominates at a certain lRMSD range. The example from Fig. 10 shows how each method returns a larger and larger motif until spanning the entire structure.

**Figure 7:**
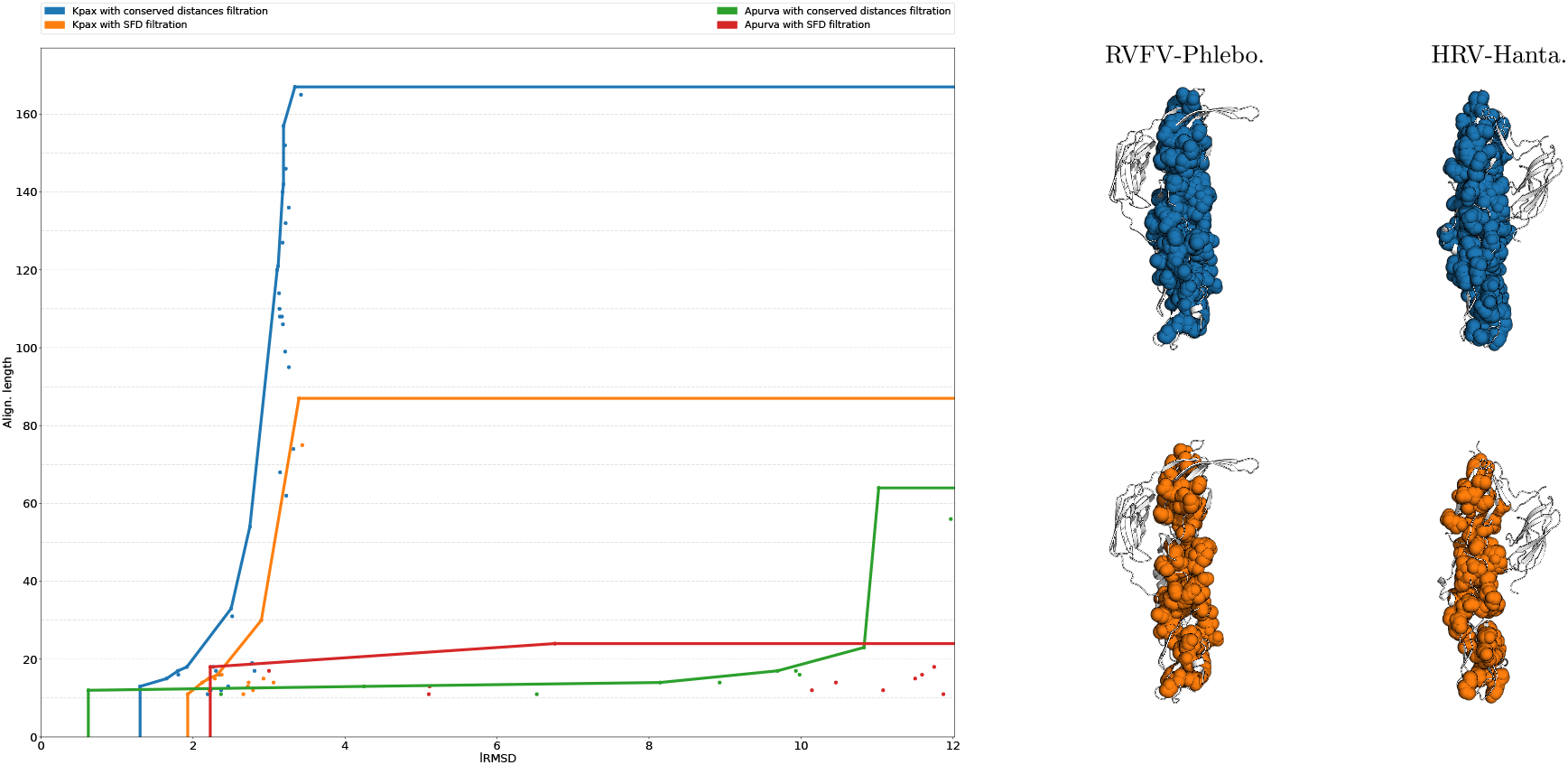
Comparison of RVFV-Phlebo. with HRV-Hanta.: (**Left**) Point clouds and Pareto envelopes of structural motifs found with each method. `Align-Kpax-CD` (blue) dominates all the other methods. (**Right**) Visualization of two structural motifs corresponding to the corner points of the `Align-Kpax-CD` (blue) and `Align-Kpax-SFD` (orange) curves.

**Figure 8:**
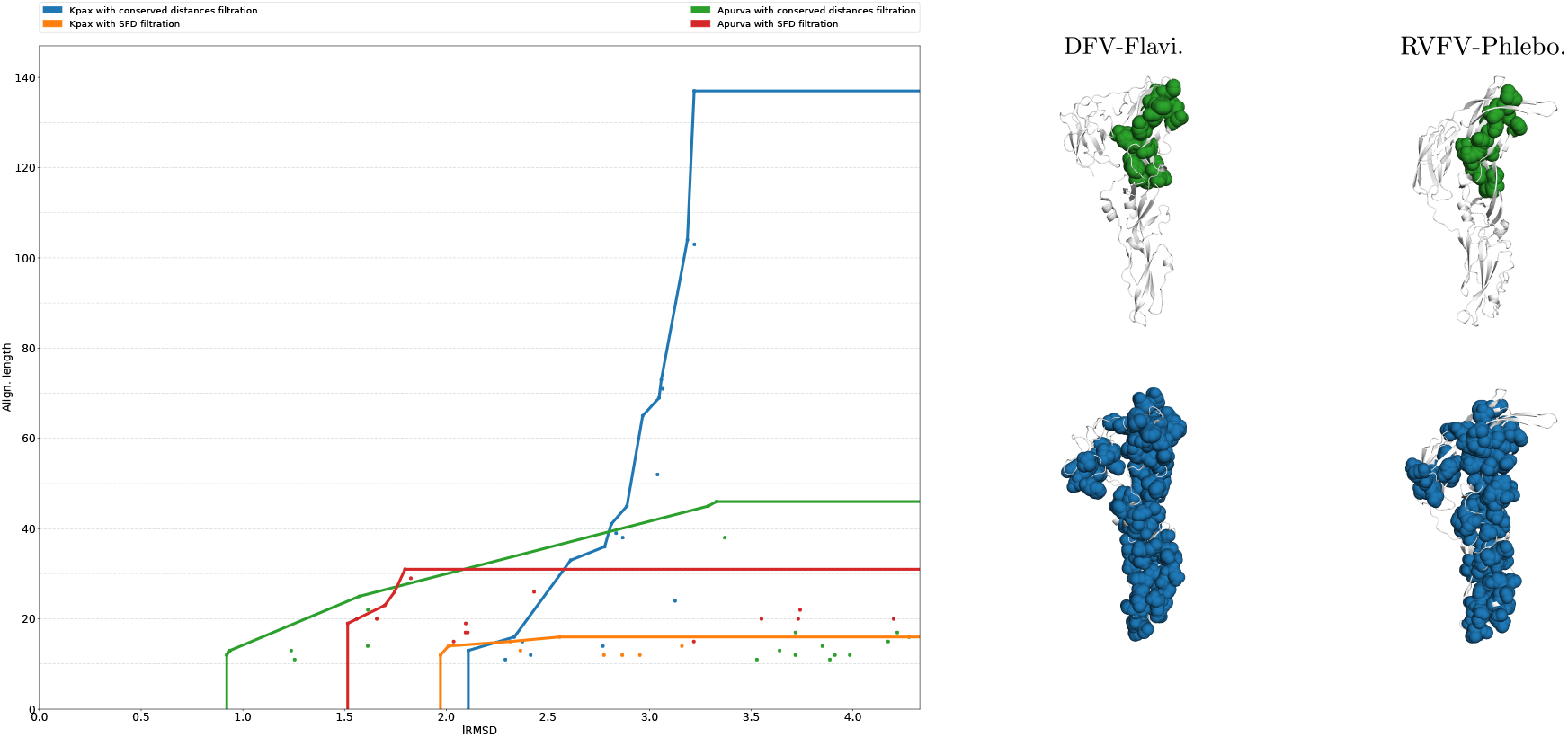
Comparison of DFV-Flavi. and RVFV-Phlebo.: (**Left**) Point clouds and Pareto envelopes of structural motifs found with each method. Initially, `Align-Apurva-CD` (Green) dominates until a critical point is reached and `Align-Kpax-CD` (Blue) takes over. (**Right**) Visualization of two structural motifs corresponding to the corner points of the `Align-Apurva-CD` (Green) and `Align-Kpax-CD` (Blue) curves.

**Figure 9:**
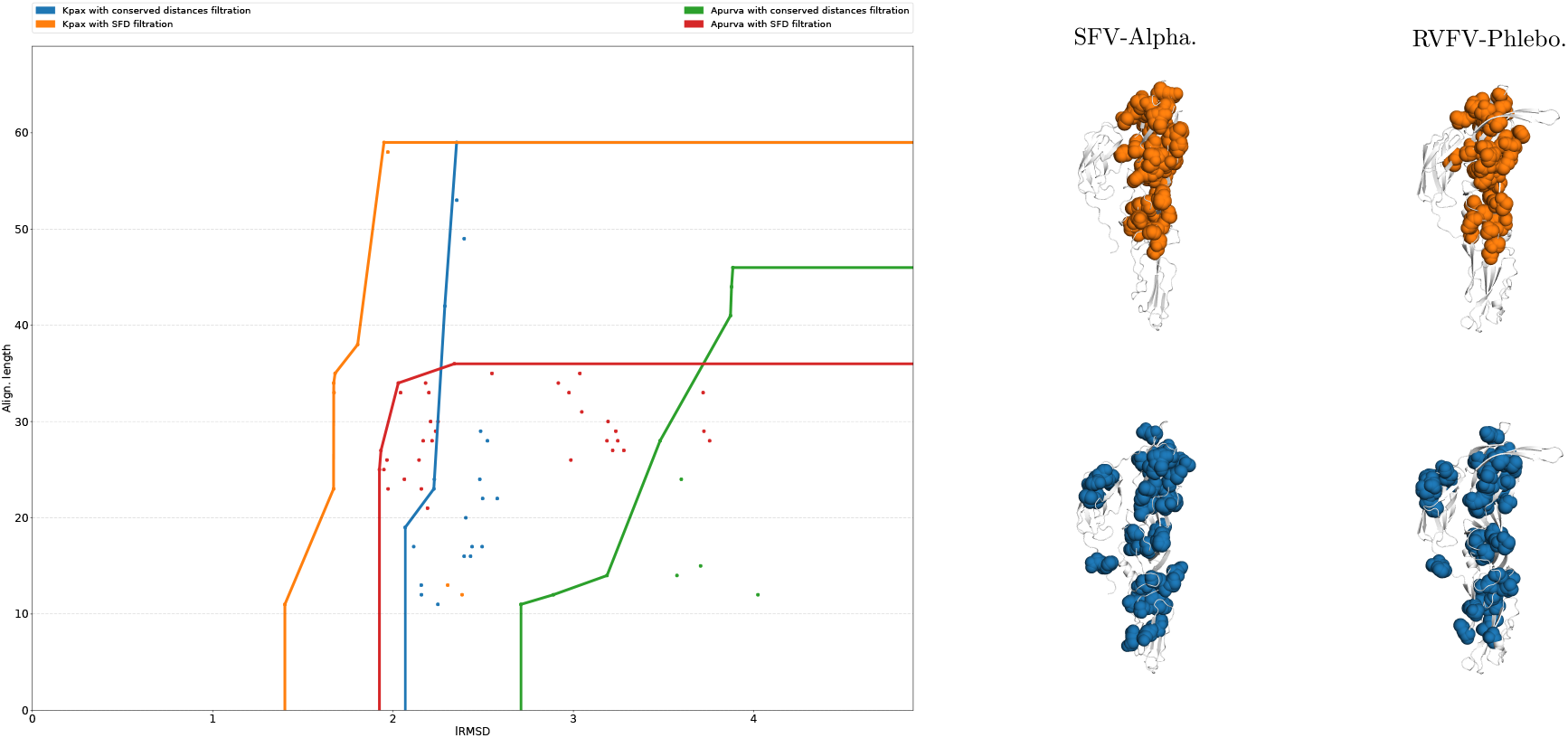
Comparison of SFV-Alpha. and RVFV-Phlebo.: (**Left**) Point clouds and Pareto envelopes of structural motifs found with each method. `Align-Kpax-SFD` (Orange) dominates all the other methods. (**Right**) Visualization of two structural motifs corresponding to the corner points of the `Align-Kpax-SFD` (Orange) and `Align-Kpax-CD` (Blue) curves.

**Figure 10:**
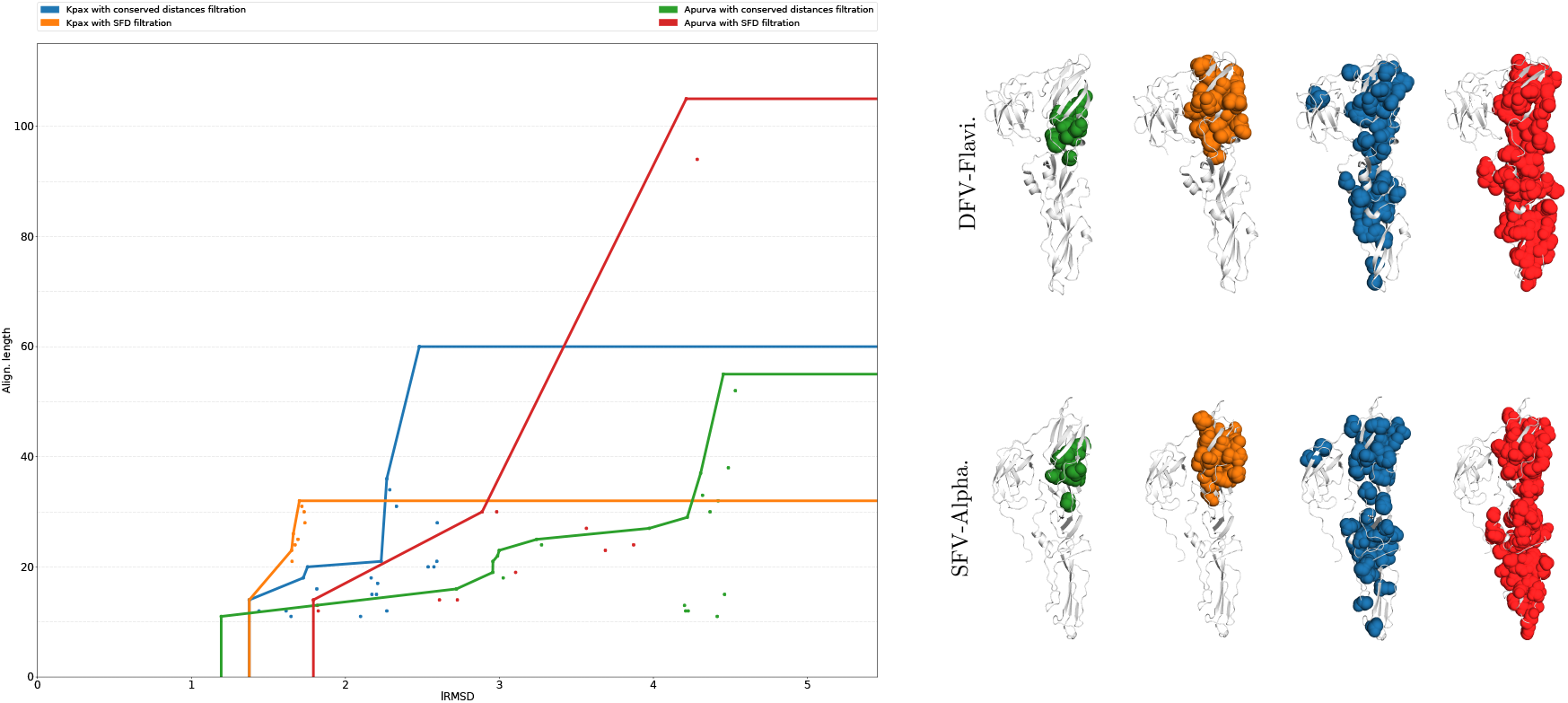
Comparison of DFV-Flavi. and SFV-Alpha.: (**Left**) Point clouds and Pareto envelopes of structural motifs found with each method. Initially, `Align-Apurva-CD` (Green) dominates until a critical point is reached and `Align-Kpax-CD` (Blue) takes over. (**Right**) Visualization of motifs corresponding to the corner points of each method.

To summarize, although `Align-Kpax-CD` dominates more often (w.r.t motif lRMSD and sizes), there is no clear better method. Qualitatively, the motifs returned by each method are quite different, the rule being that with SFD filtration motifs tend to be more localized and connected, compared to CD filtration which returns larger albeit more dispersed motifs.

### 4.4 Comparisons against flexible aligners

As noted in Introduction, our methods are not direct competitors of flexible aligners since a key goal is to capture the various structural conservation scales a seed alignment may contain.

Nevertheless, we compare our aligners against classical flexible aligners, in particular `FATCAT` and `Kpax`, using the 10 *difficult* structures from [23] (SI Section 4.4, SI Tables 4, 5, 6 and 7).

`Align-Kpax-SFD` and `Align-Apurva-SFD` fail to find motifs on certain structures. This is to be expected on smaller structures as they are prone to accretion. In most instances our methods yield RMSD_Comb_. values which are a significant improvement to the lRMSD of the `FATCAT` alignment, although at a cost in size. Recall that motifs are constrained areas of the structures. In some instances, RMSD_Comb_. is comparable to `FATCAT` (slightly better or slightly worse). In a few cases, our methods yield poor results: a striking example is that of 1CRL vs 1EDE for which we find a RMSD_Comb_. of 7.47 (against a lRMSD of 3.55 for `FATCAT`). These poor results are due to the dependency to the seed alignment and the parameterization of our method. In the same example (1CRL vs 1EDE), the seed alignment has a lRMSD of 8.08 and *τ*/ = 0.8. Running the same comparison with *τ*/ = 0.3 yields a very different result: 84 residues involved and RMSD_Comb_. = 2.38. This is a demonstration of how constraining *τ*/ enables the discovery of more conserved motifs. Seeding iterative alignments with our motifs yields results that are comparable to `FATCAT`. These alignments are very similar to the ones provided by `Kpax` in terms of lRMSD and size. However comparing structural alignments based only on the lRMSD is very constraining and these final alignments should not be dismissed, they can provide additional information.

**Remark 4** *Normally, after computing an alignment*, `Kpax` *checks for distant aligned pairs and removes them from the alignment [12]. The* `SBL` *implementation of* `Kpax` *does not do this by default, this can create some poor alignments (w.r.t lRMSD). As already noted, by running* `Kpax` (`SBL`) *on 1CRL vs 1EDE yields an alignment with a high lRMSD (8.08). By adding a distance threshold of* 0.7 *on aligned residues, we obtain a 137 residue long alignment with a lRMSD of* 1.72.

## 5 Discussion and outlook

Molecular flexibility is a continuous process, with characteristic spatial and time scales, so that a key difficulty in understand the role of flexibility and dynamics in protein function is to perform a multiscale analysis of structurally conserved motifs. Our work precisely addresses this task, by proposing a generic framework to automatically detect the multiple flexibility scales which may exist when comparing two structures, be they conformations of the same molecule, or homologous proteins. To this end, our framework bootstraps from a seed alignment, and further exploits the hierarchical information contained in so-called filtrations and the associated persistence diagrams, defined from distance difference matrices.

Our motifs naturally accommodate a hierarchical representation in terms of Hasse diagram, which provides the basis of multiscale analysis. As a first intent, terminal motifs (i.e. motifs defining roots of the aforementioned Hasse diagram) can be used in conjunction with the recently proposed combined lRMSD. More precisely, since a motif has its own optimal rigid motion and lRMSD, these lRMSD can be combined into a global measure to compare two structures. We show that the resulting combined RMSD may reduce the global lRMSD by one order of magnitude, providing a tool of unprecedented accuracy to compare two structures which otherwise may appear as radically different from a structural standpoint.

Importantly, our framework enjoys two major design choices. The first one is the seed alignment used, which may favor topological information (conserved contacts, via contact map optimization), or geometric information (e.g. conserved backbone geometry). Practically, we tested two seed aligners, namely `Apurva`, which solves the contact map overlap problem with guarantees, and `Kpax`, an iterative aligners favoring the coherence of backbone geometry. The second one is the type of filtration used, which may favor either motifs stemming from connected regions in 3D (defined via space filling diagrams), or motifs stemming from conserved distance between distant amino-acids. Each of the four resulting structural aligner targets a specific type of flexibility, and we exhibit such examples for class II fusion proteins.

In a more general perspective, these examples refer to different scenarios in terms of molecular mechanisms. The presence of structurally distant yet conserved motifs may be related to cooperative and allosteric phenomena. The presence of compact and conserved motifs, such as those observed in a classical hinge motion, may just reveal conformational changes via relative domain motions. In any case, strongly conserved motifs may hint at regions coupled to a specific function. Under this assumption, the sub-sequences associated to motifs may be used to design hybrid sequence-structure based (profile HMM) models of protein families. This strategy has been successfully pursued in a companion paper [31], where two of our methods (`Align-Kpax-CD, Align-Kpax-SFD`) identified from UniProtKB unknown class II fusion proteins in drosophilia melanogaster. Importantly, the notion of (local) motif, rather than (overall) structural alignment, turned out to be instrumental [31].

Our methods to report structural motifs are made available via the Structural Bioinformatics Library (http://sbl.inria.fr). We anticipate that they will be of interest in all problems dealing with structural analysis, and also with the identification / annotation of distant homologs.

As future work of utmost interest, we wish to point out the interest of our motifs to seed iterative structural alignments. While such alignments have traditionally focused on a small number of (optimal) solutions, in using the variety of our motifs, it might be possible to thoroughly explore the space of similar motifs.

Successfully implementing this idea would yield the counterpart of potential energy landscape exploration algorithms.

## Acknowledgments

N. Malod-Dognin is acknowledged for stimulating discussions.

## Abbreviations

a.a.: amini-acid
SFD: space filling diagram
CD: conserved distances
PD: persistence diagram

